# Transcriptional Activation of Estrogen Receptor-alpha and Estrogen Receptor-beta from Elephant Shark (*Callorhynchus milii*)

**DOI:** 10.64898/2025.12.11.693795

**Authors:** Ya Ao, Haruka Narita, Wataru Takagi, Susumu Hyodo, Michael E. Baker, Yoshinao Katsu

**Author notes:** Correspondence: Yoshinao Katsu, Michael E. Baker.

## Abstract

Humans and other vertebrates contain two estrogen receptors (ERs), ERα and ERβ. Among cartilaginous fish (sharks, rays, skates), which are chondrichthyans that evolved about 425 million years ago, only activation by steroids of ERβ orthologs has been characterized. To remedy this gap in understanding estrogen signaling in chondrichthyans, we studied estrogen activation of orthologs of human ERα and ERβ from elephant shark (*Callorhynchus milii*). Unexpectedly, we found that *C. milii* contained three estrogen-responsive ERα genes: ERα1 (596 amino acids), ERα2 (600 amino acids), and ERα3 (599 amino acids) with strong sequence similarity to each other. We also found an estrogen-unresponsive gene, ERα4 (561 amino acids), with a 39 amino acid deletion in the DNA-binding domain. An estrogen-responsive ERβ ortholog (580 amino acids) also was present in *C. milii*. The three active *C. milii* ERαs are of similar length to human ERα (595 amino acids); however, *C. milii* ERβ is longer than human ERβ (530 amino acids). We studied transcriptional activation of ERα and ERβ by estradiol (E2), the main reproductive estrogen in humans. We also studied estrone (E1), the main postmenopausal estrogen, and estriol (E3), which is synthesized during pregnancy. We determined the half-maximal response (EC50) and fold-activation to E2, E1, and E3 of *C. milii* ERα1, ERα2, ERα3, and ERβ. Among these estrogens, E2 had the lowest EC50 for all four ERs. Fold-activation by E2 and E3 was similar for ERα1, ERα2, ERα3, and ERβ. Overall, estrogen activation of *C. milii* ERα and ERβ was similar to that for human ERα and ERβ, indicating substantial conservation of the vertebrate ER during the 425 million years since the divergence of cartilaginous fish and humans from a common ancestor.

## 1. Introduction

Estrogens are a class of steroid hormones that have diverse physiological activities in females and males in humans and other vertebrates [1-5]. Estrogens act by binding to estrogen receptors (ERs) [5-6], which are transcription factors that belong to the nuclear receptor family, which also includes receptors for progestins, androgens, and corticosteroids [7-9]. To date, two distinct estrogen receptor gene: estrogen receptor alpha (ERα) and estrogen receptor beta (ERβ), respectively, have been isolated in mammals and other vertebrates [10-12]. ERα and ERβ have strong sequence similarity (97%) in their DNA binding domains, but only about 55% similarity in their ligand-binding domains.

ERα is expressed in reproductive tissues (uterus, ovary), breast, kidney and bone, while ERβ is expressed in the ovary and male <u>reproductive organs</u>, (prostate), colon, kidney and the immune system. The response of ERα and ERβ in humans and other vertebrates to 17β-estradiol (E2), the main physiological estrogen, as well as to estrone (E1) a postmenopausal estrogen, and estriol (E3), which is synthesized during pregnancy (Figure 1) has been studied extensively [5,6,10,13]. Although ERα and ERβ can have opposing actions, both receptors are activated by binding estrogens, which induce a conformational change that promotes binding to specific DNA sequences and the initiation of estrogen-dependent gene transcription. ERα promotes tissue proliferation, while ERβ may act as a suppressor of ERα-mediated proliferation.

**Figure 1.**
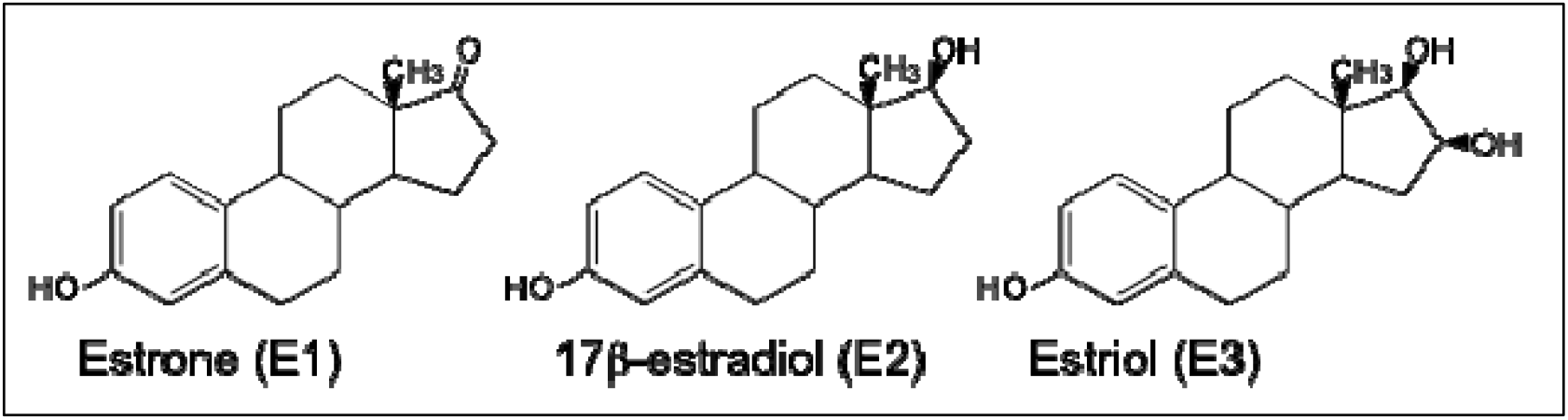
Structures of estrogen. Estrone (E1), 17β-estradiol (E2), and estriol (E3) are the natural estrogens. E1 is a minor female sex hormone and serves mainly as a precursor of E2. E3 is produced in the placenta during pregnancy. E2 is a major female sex hormone mainly produced in the ovary [38, 39].

Although ERs with sequence similarity to ERα [14,15] and ERβ [14-16] have been isolated from elasmobranchs (sharks, rays, and skates), only the response to steroids by ERβ from the cloudy catshark (*Scyliorhinus torazame*) and whale shark (*Rhincodon typus*) has been studied [16]. Estrogen activation of a gnathostome ERα has not been reported. Elasmobranchs, along with Holocephali, comprise the two extant subclasses of Chondrichthyes, jawed fishes with skeletons composed of cartilage rather than calcified bone [17]. Chondrichthyes occupy a key position relative to fish and terrestrial vertebrates, having diverged from bony vertebrates about 450 million years ago [14,18]. As the oldest group of extant jawed vertebrates (gnathostomes), the properties of ERα and ERβ in cartilaginous fishes can provide important insights into the early evolution of ERα and ERβ from an ancestral ER in lamprey [19]. Moreover, elephant shark has the slowest evolving genome of all known vertebrates, including the slowly evolving coelacanth [14], which makes elephant shark genes a good system to study the evolution of the ER, including the early evolution of selectivity of the ER in a vertebrate and the subsequent divergence in steroid selectivity in human ER, in other vertebrate ERs and in ray-finned fish ERs [20]. Another point of interest for the ER in elephant shark is that elephant sharks contain three full-length ERαs [14], raising the question of each how these three ERαs compare with each other and with mammalian ERαs in their response to estrogens.

To remedy this gap in our understanding of ERα activation in vertebrates, we have studied the response to estradiol, estrone, and estriol of three full-length *Callorhinchus milii* ERαs, as well as *C. milii* ERβ. Here we report that despite the strong sequence similarity among *C. milii* ERα1, ERα2, ERα3, and ERβ, there are some differences in their half-maximal response (EC50) and fold-activation to these three physiological estrogens. We find that E2 had the lowest EC50 for all four ERs. E2 and E3 had a similar fold activation for ERα1, ERα2, ERα3, and ERβ. Overall, estrogen activation of *C. milii* ERα and ERβ was similar to that for human ERα and ERβ [5,7], indicating substantial conservation of the vertebrate ER in the 425 million years since the divergence of cartilaginous fish and humans from a common ancestor.

## 2. Materials and methods

### 2.1. Animals

*C. milii* elephant sharks of both sexes were collected in Western Port Bay, Victoria, Australia, using recreational fishing equipment, and transported to Primary Industries Research Victoria, Queenscliff, using a fish transporter. The animals were kept in a 10-t round tank with running seawater (SW) under a natural photoperiod for at least several days before sampling. Animals were anesthetized in 0.1% (w/v) 3-amino benzoic acid ethyl ester (Sigma-Aldrich, St. Louis, MO, USA). After the decapitation of the animal, tissues were dissected out, quickly frozen in liquid nitrogen, and kept at −80°C. All animal experiments were conducted according to the Guideline for Care and Use of Animals approved by the committees of University of Tokyo as described in [21] [Animal Ethics Committee of the Ocean Research Institute of the University of Tokyo], Approval Code: [17-3], and Approval Date: [March 29, 2005]).

### 2.2. Chemical reagents

Estrone (E1), 17β-estradiol (E2), and estriol (E3) were purchased from Sigma-Aldrich Corp. (St. Louis, MO). All chemicals were dissolved in dimethyl-sulfoxide (DMSO). The concentration of DMSO in the culture medium did not exceed 0.1%.

### 2.3. Cloning of estrogen receptors

When we began this project, the genome of *C. milii* had not yet been sequenced [14]. Thus, we needed to sequence an ER in *C. milii* as a first step in determining the number of ERs and their sequences in *C. milii*. For this goal, we used two conserved amino acid regions in the DNA-binding domain (GYHYGVW) and the ligand-binding domain (NKGM/IEH) of vertebrate ERs to design the degenerate oligonucleotides to clone *C. milii* ERs. The second PCR used the first PCR amplicon, and nested primers that were selected in the DNA binding domain (CEGCKAF) and the ligand binding domain (NKGM/IEH). As a template for PCR, the first-strand cDNA was synthesized using total RNA isolated from the ovary. The amplified DNA fragments were subcloned with TA-cloning plasmids pCR2.1-TOPO (Invitrogen, Carlsbad, CA). The 5’ and 3’ ends of the ER cDNAs were amplified by rapid amplification of the cDNA end (RACE) using a SMART RACE cDNA Amplification kit (BD Biosciences Clontech., Palo Alto, CA). To amplify the isoforms of elephant shark ER, we applied a PCR-based cDNA amplification technique using the primers et (5’-TGAAGTGTGATCGTCCAGGCGACAG-3’ and 5’-CAAGCTGGAGGATAAGACATCGAC-3’). Sequencing was performed using a BigDye Terminator Cycle Sequencing kit and analyzed on the Applied Biosystems 3730 DNA Analyzer.

We used the *C. milii* extracted RNA to clone ERα1, ERα2, ERα3, ERα4, and ERβ, which had sequences corresponding to the sequences of the *C. milii* genome deposited by Venkatesh et al., 2014 [14].

### 2.4. Database and sequence analyses

All sequences generated were searched for similarity using BLASTn and BLASTp at the web servers of the National Center of Biotechnology Information (NCBI). We found four ERα genes and one ERβ gene [14], which were aligned using ClustalW [22].

### 2.5. Phylogenetic tree analysis

The evolutionary history was inferred using the Neighbor-Joining method [23]. The optimal tree is presented in this study. The percentage of replicate trees in which the associated taxa clustered together in the bootstrap test (1000 replicates) is shown next to the branches [24]. The tree is drawn to scale, with branch lengths in the same units as those of the evolutionary distances used to infer the phylogenetic tree. The evolutionary distances were computed using the Poisson correction method and are in units of the number of amino acid substitutions per site. Evolutionary analyses were conducted in MEGA 11 [25].

### 2.6. Reporter gene assay

Full-length estrogen receptors were amplified using specific forward and reverse primers designed at start and stop codons and cloned into the mammalian expression vector pcDNA3.1 (Invitrogen). In addition, ERs with a FLAG-tag added to the N-terminus were also prepared by PCR. A reporter construct, pGL4.23-4xERE was produced by subcloning of oligonucleotides having 4xERE into the *Kpn*I-*Hind*III site of pGL4.23 vector (Promega). All cloned DNA sequences were verified by sequencing.

Reporter gene assays using full-length ERs were performed in Human Embryonic Kidney 293 cells (HEK293 cells). HEK293 cells were seeded in 24-well plates at 5 x 10^4^ cells/well in phenol-red-free Dulbecco’s modified Eagle’s medium with 10% charcoal/dextran-treated fetal bovine serum. After 24 h, the cells were transfected with 400 ng of reporter construct, 25 ng of pRL-TK (as an internal control to normalize the variation in transfection efficiency; contains the *Renilla reniformis* luciferase gene with the herpes simplex virus thymidine kinase promoter), and 200 ng of pcDNA3.1-estrogen receptor using polyethylenimine (PEI). After 5 h of incubation, ligands were applied to the medium at various concentrations. After an additional 43 h, the cells were collected, and the luciferase activity of the cells was measured with the Dual-Luciferase Reporter Assay System. Promoter activity was calculated as firefly (*P. pyralis*)-luciferase activity/sea pansy (*R. reniformis*)-luciferase activity. The values shown are mean ± SEM from three separate experiments, and dose-response data, which were used to calculate the half maximal response (EC50) for each steroid, were analyzed using GraphPad Prism (Graph Pad Software, Inc., San Diego, CA). All experiments were performed in triplicate.

## 3. Results

### 3.1. Cloning and sequence analysis of cDNAs for four elephant shark ERα genes and one ERβ gene

Using standard RT-PCR and RACE techniques, we successfully cloned four full-length ERs, designated as ERα1, ERα2, ERα3 (Figure 2A, B), and ERβ (Figure 2C), from elephant shark ovary RNA. The cDNA for elephant shark ERα1 predicted a 596 amino acid protein with a calculated molecular mass of 66.6 kDa (GenBank accession no. LC068847), and the cDNA for elephant shark ERβ predicted a 580 amino acid protein with a calculated molecular mass of 64.9 kDa (GenBank accession no. LC068848), Figure 2A, B. In addition, we cloned the other two full-length elephant shark ERs: ERα2 and ERα3, which are found in GenBank (GenBank accession no. XM_007894403 for ERα2, XM_007894404 for ERα3). ERα4 (GenBank accession no. XM_007894406) with a 39 amino acid deletion in the DNA-binding domain was also cloned (Figure 2A, B).

**Figure 2.**
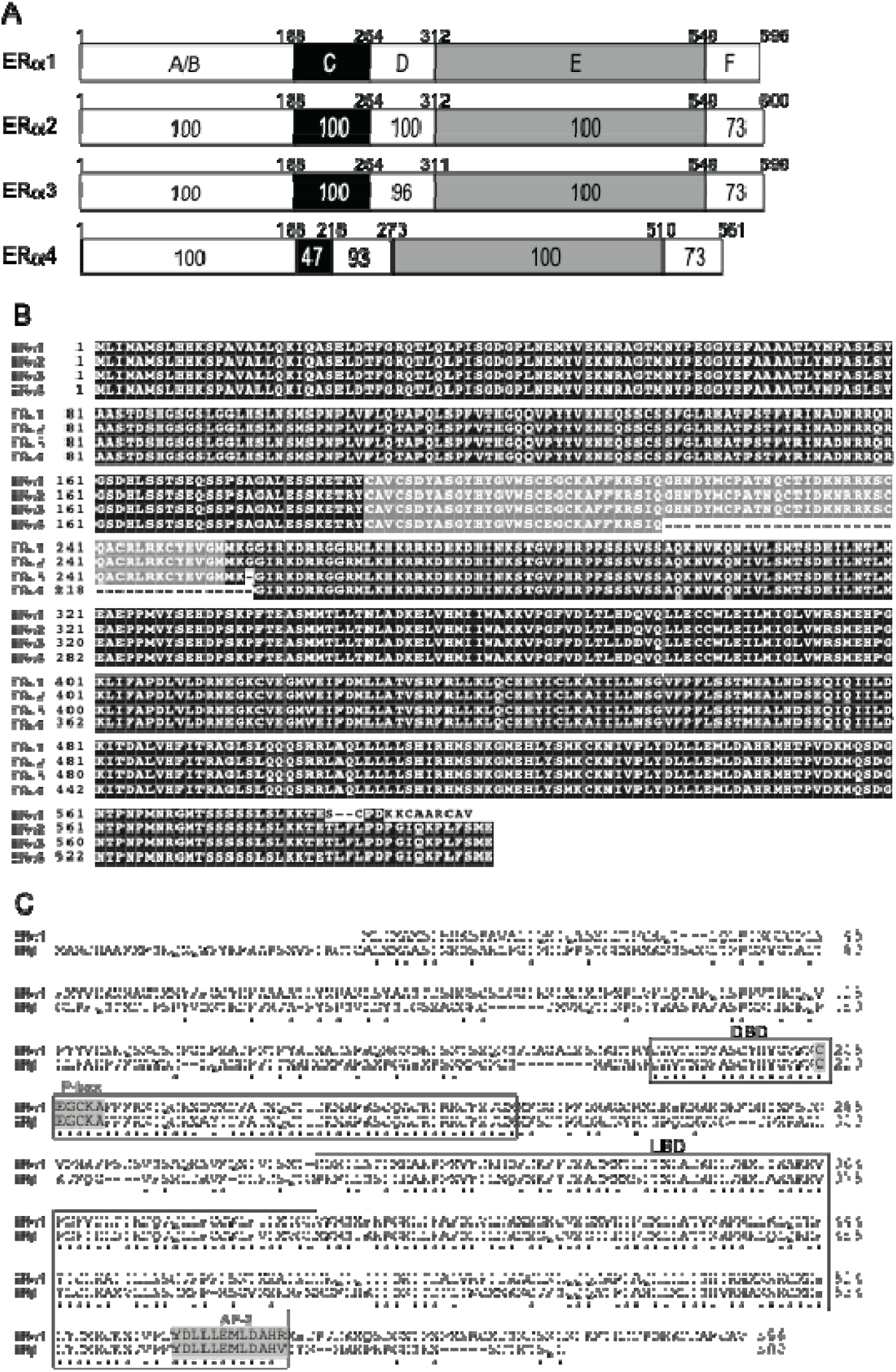
Amino acid sequences of the elephant shark ERs. (A) Comparison of functional domains of four elephant shark ERα genes. Elephant Shark ERα1, ERα2, and ERα3 show strong sequence conservation of their A/B, C, D, and E domains. ERα4 has a deletion in the DNA-binding domain [C domain], which we propose explains its lack of transcriptional activation by estradiol. (B) Sequence alignment of Elephant Shark ERα1, ERα2, ERα3, and ERα4. There is excellent conservation of the four-elephant shark ERα genes. (C) Aligned sequences of the elephant shark ERα1 and ERβ. The DBD and LBD are indicated by open boxes. Residues are important for DNA response element recognition (the P-box), and the AF-2 region, which mediates contact of the LBD with transcriptional coactivators, is shaded in gray. Numbers to the right indicate amino acid position. Asterisks indicate residues conserved in both ERα1 and ERβ.

The elephant shark ERα and ERβ sequences contain the five recognizable steroid hormone receptor sub-domains in the expected order: the N-terminal region (A/B domain), DNA binding domain, DBD (C domain), hinge region (D domain), ligand binding domain, LBD (E domain), and C-terminal extension (F domain) (Figure 2A). In Figure 2A we compare the functional domains of ERα1, ERα2, ERα3 and ERα4. Figure 2A shows that there is excellent conservation among these ERs of the A/B, C, D, E and F domains. This strong sequence conservation also is seen in the amino acid alignment of ERα1, ERα2, ERα3 and ERα4 shown in Figure 2B. Figure 2B shows the deletion in the DBD of ERα4, which probably explains the lack of transcriptional activation by estrogens of *C. milii* ERα4. The amino acid sequence of ERα4 is closest to ERα3. Overall, Figures 2A and 2B show the strong sequence conservation in the four *C. milii* ERα genes.

In Figure 2C, we show a sequence alignment of *C. milii* ERα1 and ERβ. The DNA response element recognition motif (P-box, CEGCKA) in the DBD and the AF-2 motif (core LLLEML region) in the LBD, which is required for interaction with transcriptional co-activators, are conserved in all four-elephant shark ERα sequences and in ERβ (Figure 2B and 2C).

### 3.2. Comparison of functional domains on human ERα and ERβ with functional domains in elephant shark ERα and ERβ

To gain additional insights into the evolution of the *C. milii* ERs, we compared the functional domains of elephant shark ERα1to elephant shark ERβ (Figure 3A), of elephant shark ERα1 to human ERα and human ERβ (Figure 3C), and of elephant shark ERβ to human ERα and human ERβ (Figure 3D). We also compared human ERα and ERβ to each other (Figure 3B).

**Figure 3.**
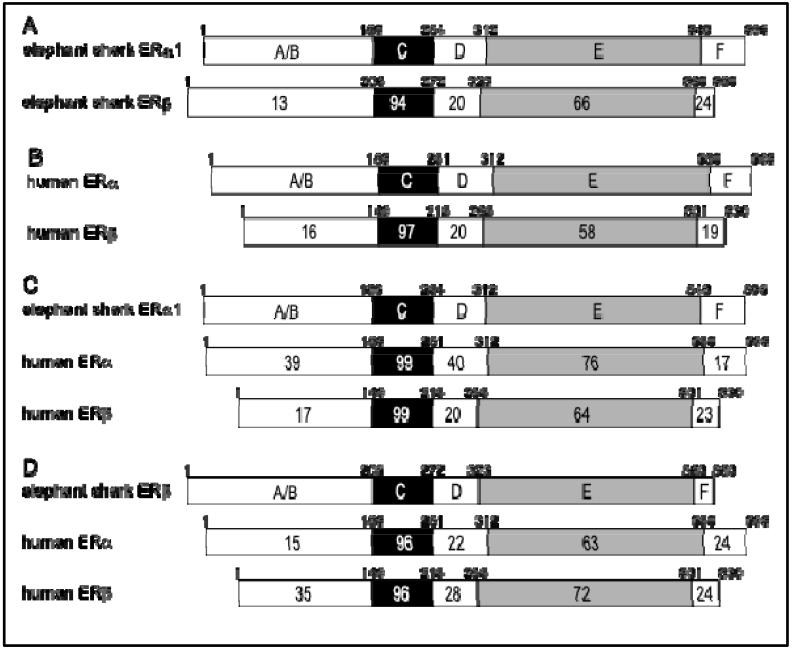
Comparison of the functional domains of elephant shark ERs with human ERs. The functional domains A/B through F are shown schematically with the number of amino acid residues indicated. The percentage of amino acid identity is shown. (A) Comparison of elephant shark ERα1 with elephant shark ERβ. The N-terminal (A/B) domains of elephant shark ERα1 and ERβ show strong divergence. There is also some divergence between the ligand (E) binding domain on elephant shark ERα1 and ERβ. (B) Comparison of human ERα (accession no. NM000125) with human ERβ (accession no. AB006590). The domains of human ERα with human ERβ have similar sequence similarity to each other as elephant shark ERα1 and ERβ. (C) Comparison of elephant shark ERα1 with human ERα and human ERβ. Elephant shark ERα1 is closer to human ERα than to human ERβ. (D) Comparison of elephant shark ERβ with human ERα and human ERβ. Elephant shark ERβ is closer to human ERβ than to human ERα.

Figure 3A shows the excellent conservation of the DNA-binding domains [C domains] (94% identity) and ligand-binding domains [E domains] (66% identity) in elephant shark ERα1 and ERβ, in contrast to the weak conservation of the A/B domains (13% identity) at their N-terminus. Comparison of Figures 3A and B reveals that the DBD in human ERα and *C. milii* ERα1 and ERβ have similar sequence conservation, while the LBD sequence in *C. milii* ERα1 and *C. milii* ERβ are more conserved (66% identity) than the LBD in human ERα and ERβ (58% identity).

Figure 3C shows the strong sequence conservation of the DBD in elephant shark ERα1 and human ERα and ERβ. Figure 3C also shows that there is stronger conservation (76%) of the LBD in elephant shark ERα1 and human ERα, compared to the similarity (64%) of the corresponding LBD in elephant shark ERα1 to human ERβ. Figure 3D shows the strong sequence conservation of the DBD in elephant shark ERα1 and human ERα and ERβ. Figure 3D also shows the stronger conservation (72%) of the LBD in elephant shark ERβ and human ERβ, compared to the similarity (64%) to the corresponding LBD in human ERα.

### 3.3. Comparison of functional domains on elephant shark ERα and ERβ with functional domains in zebrafish, and whale shark

To gain an additional insight into the evolution of elephant shark ERα and ERβ, we compared their sequences to ERα and ERβ sequences in zebrafish and whale shark (Figure 4).

**Figure 4.**
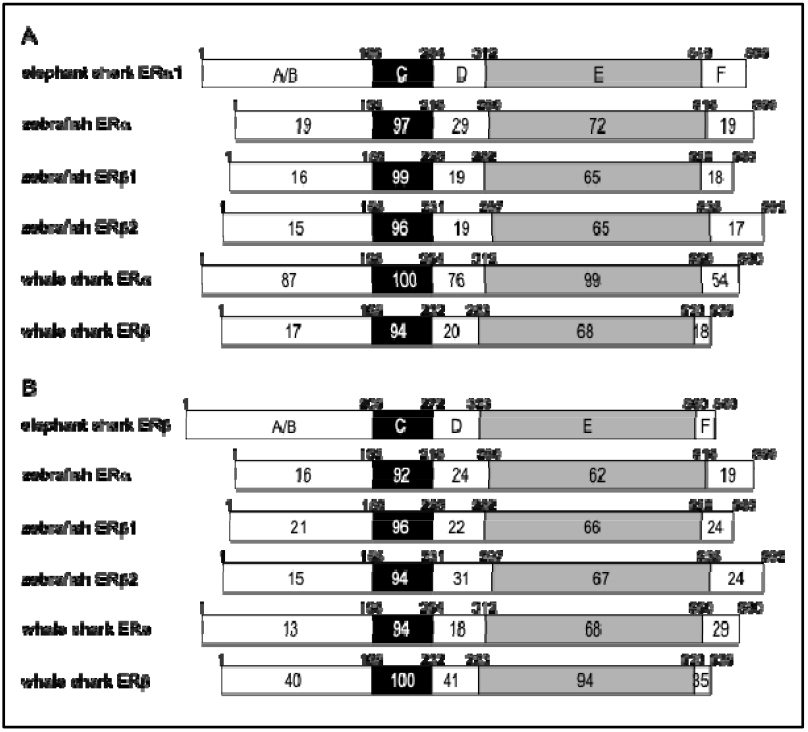
Comparison of the functional domains of elephant shark ERs with ERs from zebrafish and whale shark. The functional domains A/B through F are shown schematically with the number of amino acid residues indicated. The percentage of amino acid identity is shown. (A) Comparison of elephant shark ERα1 with zebrafish and whale shark ERs. (B) Comparison of elephant shark ERβ with zebrafish and whale shark ERs. Genbank accession no. NM_152959 (zebrafish ERα), AF516874 (zebrafish ERβ1), AF349413 (zebrafish ERβ2), XM_048600891 (whale shark ERα), and AB551716 (whale shark ERβ).

The sequence comparisons (Figure 4) between elephant shark ERα and ERβ domains with ERα and ERβ domains in whale shark revealed an unexpected sequence similarity (87% identity) between the A/B domain on elephant shark ERα and whale shark ERα. The A/B domain in whale shark ERβ and elephant shark ERβ has 40% identity. Interestingly, the A/B domains on elephant shark ERα and ERβ domains have 39% and 35% sequence identity, respectively, with the A/B domains on human ERα and human ERβ, respectively.

The stronger conservation of the LBD in elephant shark ERβ and human ERβ (72% identity), compared to the similarity to the corresponding LBD in human ERα (63% identity) is shown in Figure 3C. The LBD of elephant shark ERα1 is closer to the LBD in human ERα than the LBD in human ERβ, and the LBD of elephant shark ERβ is closer to the LBD in human ERβ than the LBD in human ERα (Figure 3C).

### 3.4. Phylogenetic analysis of estrogen receptors in elephant sharks, humans and other vertebrates

To better understand the relationship between elephant shark ERs and other vertebrates ERs, we constructed a phylogeny of ERs from sharks, humans and other vertebrates [Figure 5] using the Neighbor-Joining method [23]. The phylogeny in shows the early divergence of shark ERs from other ERs.

**Figure 5.**
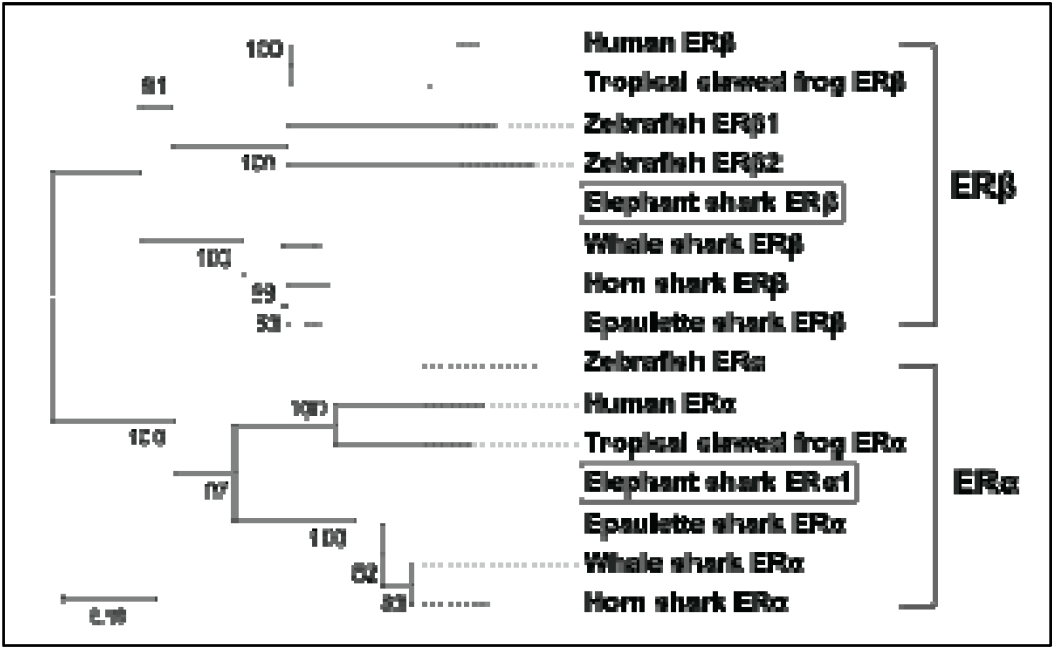
Phylogenetic analysis of estrogen receptors in elephant sharks, humans and other vertebrates. The phylogenetic tree was constructed using the maximum likelihood analysis [24]. The two elephant shark ERs are marked with open boxes. The scale bar indicates 0.1 expected amino acid substitutions per site. The GenBank accession numbers are NM_000125 (human ERα), AB006590 (human ERβ), NM_203535 (tropical clawed frog ERα), NM_001040012 (tropical clawed frog ERβ), XM_048600891 (whale shark ERα), AB551716 (whale shark ERβ), XM_068044449 (horn shark ERα), XM_068038484 (horn shark ERβ), XM_060831156 (epaulette shark ERα), XM_060829508 (epaulette shark ERβ), NM_152959 (zebrafish ERα), AF516874 (zebrafish ERβ1), AF349413 (zebrafish ERβ2).

### 3.5. Transcriptional activities of elephant shark ERα and ERβ

A transactivation assay was used to examine the response to the physiological estrogens E1, E2, and E3 of the three active isoforms of elephant shark ERα. The three isoforms, ERα showed ligand-dependent transactivation (Figure 6A, B, C). The EC50s for transcriptional activation of ERα1, ERα2, and ERα3 were 0.49 nM, 0.29 nM, and 0.25 nM (E1), 0.0059 nM, 0.016 nM, and 0.0093 nM (E2), and 0.2 nM, 0.31 nM, and 0.29 nM (E3), respectively (Table 1).

**Table 1.** Gene transcriptional activities of elephant shark ERs by estrogens in HEK293.

|  |  | E1 | E2 | E3 |
| --- | --- | --- | --- | --- |
| ERα1 | EC50 (nM) | 0.49 | 0.0059 | 0.2 |
| | Fold-Activation (± SEM) * | 1.62 (± 0.16) | 1.66 (± 0.04) | 1.57 (± 0.09) |
|  | Ratio** | 0.98 | 1.00 | 0.95 |
| ERα2 | EC50 (nM) | 0.29 | 0.016 | 0.31 |
| | Fold-Activation (± SEM) * | 1.51 (± 0.12) | 1.79 (± 0.12) | 1.82 (± 0.13) |
|  | Ratio** | 0.84 | 1.00 | 1.02 |
| ERα3 | EC50 (nM) | 0.25 | 0.0093 | 0.29 |
| | Fold-Activation (± SEM) * | 1.55 (± 0.11) | 1.75 (± 0.08) | 1.87 (± 0.15) |
|  | Ratio** | 0.89 | 1.00 | 1.07 |
| ERβ | EC50 (nM) | 0.14 | 0.001 | 0.02 |
| | Fold-Activation (± SEM) * | 2.87 (± 0.16) | 3.23 (± 0.26) | 3.49 (± 0.26) |
|  | Ratio** | 0.89 | 1.00 | 1.08 |

\*Fold activation values were calculated based on the 10 nM values.
\*\*Ratio values were calculated by dividing the fold-activation value of estrogens by the value of E2.

**Figure 6.**
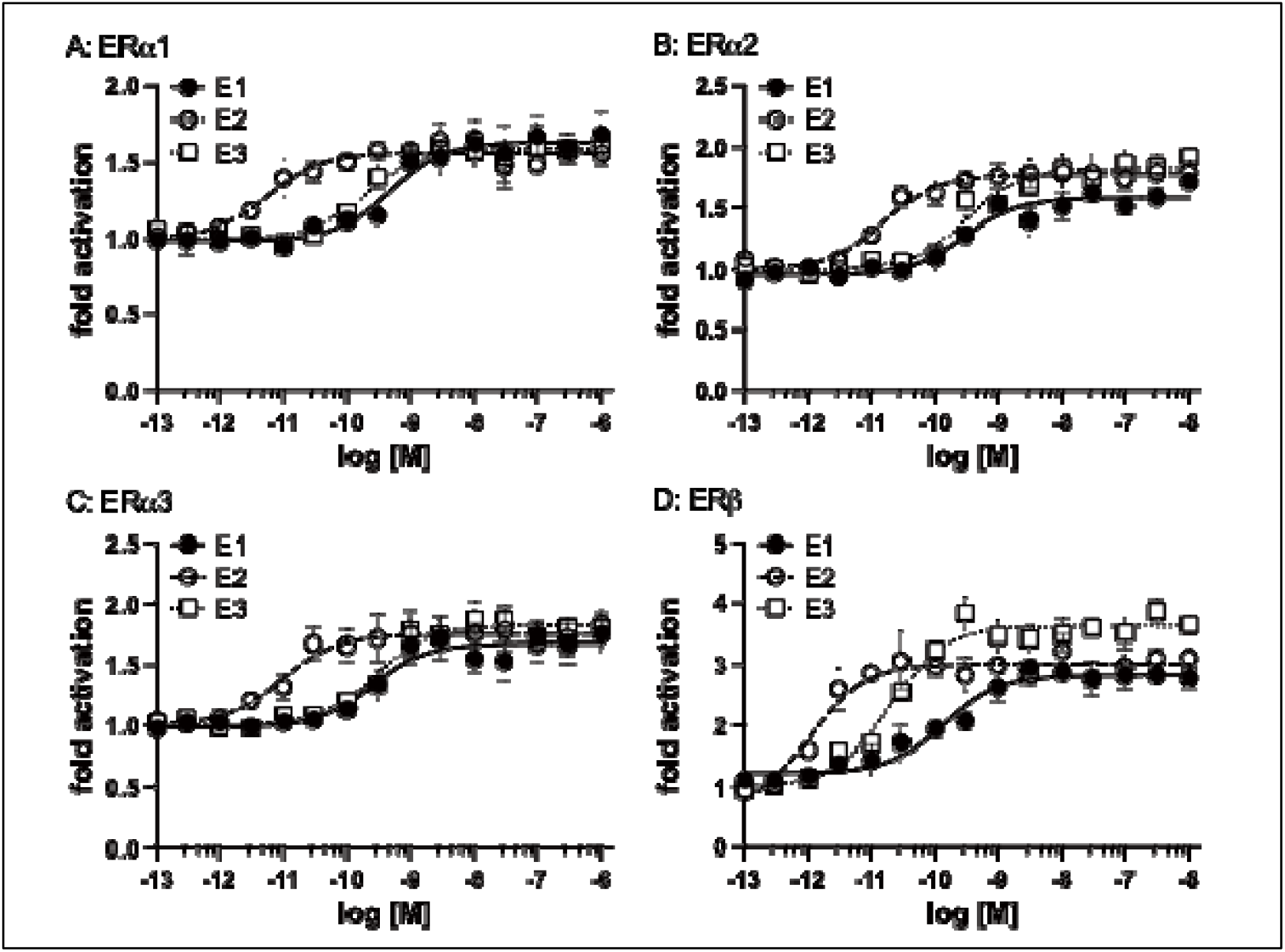
Transcriptional activation of elephant shark ERs. Elephant shark ERs were transfected into HEK293 cells with an ERE-driven reporter gene. Concentration-response profile for ERα1 (A), ERα2 (B), ERα3 (C), and ERβ (D) for E1, E2, and E3 (10^-13^M to 10^-6^M). Data are expressed as a ratio of a vehicle (DMSO) to other test chemicals. Each point represents the mean of triplicate determinations, and vertical bars represent the mean ± SEM.

We also examined the induction of elephant shark ERβ transcriptional activity by different concentrations of estrogens (E1, E2, and E3) (Figure 6). All three estrogens activated transcription of elephant shark ERβ in a dose-dependent manner (Figure 6D). The half-maximal response (EC50) for transcriptional activation of elephant shark ERβ was 0.14 nM for E1, 0.001 nM for E2, and 0.02 nM for E3 (Table 1). Although there was no significant difference in EC50 values for activation by an estrogen of ERα1 and ERβ, the fold activation values for E2 were higher than those for E1 and E2 (Table 1). Thus, the elephant shark estrogen receptors exhibited a ligand responsiveness observed human estrogen receptors [1-10].

## 4. Discussion

Significant advances have been made in elucidating the evolution of estrogen signaling in animals [1, 2, 9, 11, 19, 26, 27]. Amphioxus, is a chordate that contains an ER and a steroid receptor (SR) that is an ortholog of receptors for 3-keto-steroids, including testosterone, progesterone, cortisol, and aldosterone [11, 28]. Contrary to expectations, the amphioxus ER does not bind E2 or other steroids [11, 27, 28], while E2 and E1 are transcriptional activators of the SR [11, 28]. Atlantic sea lamprey contains an ER [19] that is activated by E2 and E1 [11, 27].

The estrogen receptors in Chondrichthyes, cartilaginous fishes with jaws, are basal to lobe-finned fishes and are ancestors of terrestrial vertebrates [14]. These ERs are still not fully characterized [15, 29]. The identification of ERα1, ERα2, ERα3, ERα4, and ERβ from the sequencing of the elephant shark [14] provided an opportunity to investigate estrogen signaling in a basal jawless vertebrate that evolved about 425 million years ago. We investigated the response to E1, E2, and E3 of ERα1, ERα2, ERα3, and ERβ from elephant shark expressed in HEK293 cells (Figure 5). These studies revealed that elephant shark ERα1, ERα2, ERα3, and ERβ are activated by E1, E2, and E3. We note that we found that ERα4 did not respond to an estrogen (data not shown), which we propose is due to a sequence deletion in the DBD. Thus, we did not continue studies of estrogen activation of ERα4.

Our data with estrogen activation of ERα1, ERα2, ERα3, and ERβ align with the established ligand binding characteristics of ERs in other vertebrates, thereby substantiating the conserved nature of estrogen signaling pathways [28, 30-32]. Estrogen-dependent transcriptional activity of the ERs from various species, including other sharks, revealed that the ERs from all species have a stronger response to E2 than to E3 [16, 33-37].

Instead of only one ERα gene, elephant shark contains three ERα genes with very similar sequences, which is unexpected. The strong sequence conservation of ERα4, which is not lig- and-activated, also is unexpected. Further research is needed to gain a more comprehensive understanding of the physiological function of estrogen and the multiple estrogen receptors in elephant sharks. This data should provide clues for understanding the physiological function(s) of each isoform.

In summary, this is the first report with evidence for the conservation of elephant shark ERα and ERβ compared to human ERα and ERβ, which provides a valuable perspective on the early evolutionary history of these critical nuclear receptors in vertebrates. The findings from this study not only confirm the conserved nature of ER-mediated estrogen signaling across vertebrates but also open new avenues for research into the functional diversification of these receptors across vertebrates. These findings furnish researchers with critical molecular data to examine the role of ERs in future studies, such as those examining gonadal development, reproductive biology, and developmental and evolutionary endocrinology.

## Author Contribution

**Y.A**.: Writing – review & editing, Writing – original draft, Methodology, Investigation, Formal analysis, and Data curation. **H.N**.: Writing – original draft, Methodology, Investigation, Formal analysis, Data curation. **W. T**.: Resources. **S.H**.: Resources. **M.E.B**.: Writing – review & editing, Project administration, Validation, Supervision, Conceptualization. **Y.K**.: Writing – review & editing, Project administration, Visualization, Validation, Funding acquisition, Supervision, Conceptualization.

## Data availability

All data is freely available on request to Dr. Katsu or to Dr. Baker.

## Acknowledgment

The authors are grateful to Dr. Kaori Oka for her suggestions and helpful advice. We also thank colleagues in our laboratories.

## References

[1] Bondesson M, Hao R, Lin CY, Williams C, Gustafsson JÅ. 2015. Estrogen receptor signaling during vertebrate development. Biochim Biophys Acta 1849(2), 142–151. doi: 10.1016/j.bbagrm.2014.06.005.

[2] Gao H, Dahlman-Wright K. 2011. The gene regulatory networks controlled by estrogens. Mol Cell Endocrinol. 334(1-2), 83–90. doi: 10.1016/j.mce.2010.09.002.

[3] Deroo BJ, Korach KS. Estrogen receptors and human disease. J Clin Invest. 2006 Mar;116(3):561–70. doi: 10.1172/JCI27987.

[4] Hamilton KJ, Hewitt SC, Arao Y & Korach KS 2017 Estrogen Hormone Biology. Curr Top Dev Biol 125 109–146.

[5] Paterni I, Granchi C, Katzenellenbogen JA & Minutolo F 2014 Estrogen receptors alpha (ERalpha) and beta (ERbeta): subtype-selective ligands and clinical potential. Steroids 90 13–29

[6] Heldring N, Pawson T, McDonnell D, Treuter E, Gustafsson JA, Pike AC. Structural insights into corepressor recognition by antagonist-bound estrogen receptors. J Biol Chem. 2007 Apr 6;282(14):10449–55. doi: 10.1074/jbc.M611424200.

[7] Dahlman-Wright K, Cavailles V, Fuqua SA, Jordan VC, Katzenellenbogen JA, Korach KS, Maggi A, Muramatsu M, Parker MG, Gustafsson JA. 2006. International Union of Pharmacology. LXIV. Estrogen receptors. Pharmacol Rev. 58(4), 773–781. doi: 10.1124/pr.58.4.8.

[8] Markov GV, Tavares, R, Dauphin-Villemant C, Demeneix BA, Baker ME, Laudet V. 2009. Independent elaboration of steroid hormone signaling pathways in metazoans. Proc Natl Acad Sci USA. 106(29), 11913–11918. doi: 10.1073/pnas.0812138106.

[9] Baker ME. 2011. Origin and diversification of steroids: co-evolution of enzymes and nuclear receptors. Mol Cell Endocrinol. 334(1-2), 14–20. doi: 10.1016/j.mce.2010.07.013.

[10] Kuiper GG, Enmark E, Pelto-Huikko M, Nilsson S, Gustafsson JA. Cloning of a novel receptor expressed in rat prostate and ovary. Proc Natl Acad Sci U S A. 1996 Jun 11;93(12):5925–30. doi: 10.1073/pnas.93.12.5925.

[11] Bridgham JT, Brown JE, Rodriguez-Mari A, Catchen JM, Thornton JW. 2008. Evolution of a new function by degenerative mutation in cephalochordate steroid receptors. PLoS Genet. 4(9), e1000191. doi: 10.1371/journal.pgen.1000191.

[12] Baker ME, Nelson DR, Studer RA. 2015. Origin of the response to adrenal and sex steroids: Roles of promiscuity and co-evolution of enzymes and steroid receptors. J. Steroid Biochem. Mol Biol 151, 12–24. doi: 10.1016/j.jsbmb.2014.10.020.

[13] Escande A, Servant N, Rabenoelina F, Auzou G, Kloosterboer H, Cavaillès V, Balaguer P, Maudelonde T. 2009. Regulation of activities of steroid hormone receptors by tibolone and its primary metabolites. J Steroid Biochem Mol Biol. 116(1-2):8–14. doi: 10.1016/j.jsbmb.2009.03.008.

[14] Venkatesh B, Lee AP, Ravi V, Maurya AK, Lian MM, Swann JB, Ohta Y, Flajnik MF, Sutoh Y, Kasahara M, Hoon S, Gangu V, Roy SW, Irimia M, Korzh V, Kondrychyn I, Lim ZW, Tay BH, Tohari S, Kong KW, Ho S, Lorente-Galdos B, Quilez J, Marques-Bonet T, Raney BJ, Ingham PW, Tay A, Hillier LW, Minx P, Boehm T, Wilson RK, Brenner S, Warren WC. 2014. Elephant shark genome provides unique insights into gnathostome evolution. Nature 505(7482), 174–179. doi: 10.1038/nature12826.

[15] Filowitz GL, Rajakumar R, O’Shaughnessy KL, Cohn MJ. 2018. Cartilaginous Fishes Provide Insights into the Origin, Diversification, and Sexually Dimorphic Expression of Vertebrate Estrogen Receptor Genes. Mol Biol Evol. 35(11), 2695–2701. doi: 10.1093/molbev/msy165.

[16] Katsu Y, Kohno S, Narita, H, Urushitani H, Yamane K, Hara A, Clauss TM, Walsh MT, Miyagawa S, Guillette LJ, Iguchi T. 2010a. Cloning and functional characterization of Chondrichtyes, cloudy catshark, Scyliorhinus torazame and whale shark, Rhincodon typus estrogen receptors. Gen Comp Endocrinol. 168(3), 496–504. doi: 10.1016/j.ygcen.2010.06.010.

[17] Inoue JG, Miya M, Lam K, Tay BH, Danks JA, Bell J, Walker TI, Venkatesh B. 2010. Evolutionary origin and phylogeny of the modern holocephalans (Chondrichthyes: Chimaeriformes): a mitogenomic perspective. Mol Biol Evol. 27(11):2576–2586. doi: 10.1093/molbev/msq147.

[18] Yu WP, Rajasegaran V, Yew K, Loh WL, Tay BH, Amemiya CT, Brenner S, Venkatesh B. 2008. Elephant shark sequence reveals unique insights into the evolutionary history of verte-brate genes: A comparative analysis of the protocadherin cluster. Proc Natl Acad Sci U S A. 105(10):3819–3824. doi: 10.1073/pnas.0800398105.

[19] Thornton JW. 2001. Evolution of vertebrate steroid receptors from an ancestral estrogen receptor by ligand exploitation and serial genome expansions. Proc Natl Acad Sci USA. 98(10), 5671–5676. doi: 10.1073/pnas.091553298.

[20] Baker ME. 2019. Steroid receptors and vertebrate evolution. Mol Cell Endocrinol. 496:110526. doi: 10.1016/j.mce.2019.110526.

[21] Kakumura K, Watanabe S, Bell JD, Donald JA, Toop T, Kaneko T, Hyodo S. 2009. Multiple urea transporter proteins in the kidney of holocephalan elephant fish (Callorhinchus milii). Comp Biochem Physiol B Biochem Mol Biol. 154(2), 239–247. doi: 10.1016/j.cbpb.2009.06.009.

[22] Thompson JD, Higgins DG, Gibson TJ. 1994. CLUSTAL W: improving the sensitivity of progressive multiple sequence alignment through sequence weighting, position-specific gap penalties and weight matrix choice. Nucleic Acids Res. 22(22), 4673–4680. doi: 10.1093/nar/22.22.4673.

[23] Saitou N, Nei M. The neighbor-joining method: a new method for reconstructing phylogenetic trees. Mol Biol Evol. 1987 Jul;4(4):406–25. doi: 10.1093/oxfordjournals.molbev.a040454.

[24] Felsenstein J. CONFIDENCE LIMITS ON PHYLOGENIES: AN APPROACH USING THE BOOTSTRAP. Evolution. 1985 Jul;39(4):783–791. doi:10.1111/j.1558-5646.1985.tb00420.x.

[25] Tamura K, Stecher G, Kumar S. MEGA11: Molecular Evolutionary Genetics Analysis Version 11. Mol Biol Evol. 2021 Jun 25;38(7):3022–3027. doi: 10.1093/molbev/msab120.

[26] Cotnoir-White D, Laperriere D, Mader S. 2011. Evolution of the repertoire of nuclear receptor binding sites in genomes. Mol Cell Endocrinol. 334(1-2), 76-82. doi: 10.1016/j.mce.2010.10.021.

[27] Paris M, Pettersson K, Schubert M, Bertrand S, Pongratz I, Escriva H, Laudet V. 2008. An amphioxus orthologue of the estrogen receptor that does not bind estradiol: insights into estrogen receptor evolution. BMC Evol Biol. 8, 219. doi: 10.1186/1471-2148-8-219.

[28] Katsu Y, Kubokawa K, Urushitani H, Iguchi T. 2010. Estrogen-dependent transactivation of amphioxus steroid hormone receptor via both estrogen and androgen response elements. Endocrinology 151(2), 639–648. doi: 10.1210/en.2009-0766.

[29] Awruch CA. 2013. Reproductive endocrinology in chondrichtyans: the present and the future. Gen Comp Endocrinol. 192, 60–70. doi: 10.1016/j.ygcen.2013.05.021.

[30] Green S, Walter P, Kumar V, Krust A, Bornert JM, Argos P, Chambon P. 1986. Human oestrogen receptor cDNA: sequence, expression and homology to v-erb-A. Nature. 320(6058), 134–139. doi: 10.1038/320134a0.

[31] Matthews J, Gustafsson JA. 2003. Estrogen signaling: a subtle balance between ER alpha and ER beta. Mol Interv. 3(5), 281–292. doi: 10.1124/mi.3.5.281.

[32] Katzenellenbogen BS, Katzenellenbogen JA. 2000. Estrogen receptor transcription and transactivation: Estrogen receptor alpha and estrogen receptor beta: regulation by selective estrogen receptor modulators and importance in breast cancer. Breast Cancer Res. 2(5), 335–344. doi: 10.1186/bcr78.

[33] Katsu Y, Lange A, Urushitani H, Ichikawa R, Paul GC, Cahill LL, Jobling S, Tyler CR, Iguchi T. 2007. Functional associations between two estrogen receptors, environmental estrogens and sexual disruption in the roach (Rutilus rutilus). Environ Sci Tech. 41(9), 3368–3374. doi: 10.1021/es062797I

[34] Katsu Y, Kohno S, Hyodo S, Ijiri S, Adachi S, Hara A, Guillette LJ, Iguchi T. 2008. Molecular cloning, characterization and evolutionary analysis of estrogen receptors from phylogenetically ancient fish. Endocrinology 149(12), 6300–6310. doi: 10.1210/en.2008-0670

[35] Katsu Y, Taniguchi E, Urushitani H, Miyagawa S, Takase M, Kubokawa K, Tooi O, Oka T, Santo N, Myburgh J, Matsuno A, Iguchi T. 2010c. Molecular cloning and characterization of ligand- and species-specificity of amphibian estrogen receptors. Gen Comp Endocrinol. 168(2), 220–230. doi: 10.1016/j.ygcen.2010.01.002

[36] Lange A, Katsu Y, Miyagawa S, Ogino Y, Urushitani H, Kobayashi T, Hirai T, Shears JA, Nagae M, Yamamoto J, Ohnishi Y, Oka T, Tatarazako N, Ohta Y, Tyler CR, Iguchi T. 2012. Comparative responsiveness to natural and synthetic estrogens of fish species commonly used in the laboratory and field monitoring. Aquat Toxicol 109, 250–258. doi: 10.1016/j.aquatox.2011.09.004.

[37] Yatsu R, Katsu Y, Kohno S, Mizutani T, Ogino Y, Ohta Y, Myburgh J, van Wyk JH, Guillette LJ Jr, Miyagawa S, Iguchi T. 2016. Characterization of evolutionary trend in squamate estrogen receptor sensitivity. Gen Comp Endocrinol. 238, 88–95. doi: 10.1016/j.ygcen.2016.04.005.

[38] Kuhl H. Pharmacology of estrogens and progestogens: influence of different routes of admin-istration. Climacteric. 2005 Aug;8 Suppl 1:3–63. doi: 10.1080/13697130500148875.

[39] Stanczyk FZ. 2024. Metabolism of endogenous and exogenous estrogens in women. J Steroid Biochem Mol Biol. 2024 Sep;242:106539. doi: 10.1016/j.jsbmb.2024.106539.

